# The applicability of Regional Red List assessments for soil invertebrates: First assessment of five native earthworm species in Canada

**DOI:** 10.1101/2024.08.23.608777

**Authors:** Helen R. P. Phillips, George G. Brown, Sam W. James, Jérôme Mathieu, John Warren Reynolds, Maheshi E. Dharmasiri, Claire L. Singer, Maria J. I. Briones, Heather C. Proctor, Erin K. Cameron

## Abstract

Earthworms (Annelida: Clitellata: Crassiclitellata) are prominent members of the soil community, important to many ecosystem functions. Despite this, and like many other soil invertebrates, they are rarely considered in conservation assessments, including the IUCN Red List assessments used to assess species’ extinction risk. To investigate the applicability of the IUCN Regional Red Listing protocol to soil invertebrates, we assessed the conservation status of five earthworm species known to be native to Canada using this protocol and all available occurrence records. In Canada, no earthworm species have yet been assessed by the Committee on the Status of Endangered Wildlife in Canada (COSEWIC). Due to the lack of data on population sizes and their trends, all five species were assessed using their Extent of Occurrence (EOO) (Criterion B). One species was assessed as Vulnerable, two were assessed in non-threatened categories, and two were assessed as Data Deficient. For the majority, the main threats identified were the continuing loss of potential habitat due to land conversion and resource exploitation, as well as the effects of climate change. Increasing the amount of data, including but not limited to distribution and habitat preferences, would make the assessment process easier and status decisions better supported. By undertaking regional assessments for five native earthworm species in Canada, we show that Regional Red List assessments are feasible for soil invertebrates.

## Introduction

It has long been estimated that up to 95% of all species on earth are invertebrates (Wilson 1987). Furthermore, not only is there huge diversity of organisms within soil ecosystems (up to 59% of all species on earth; Anthony et al. 2023), these organisms play key roles in many ecosystem functions and services (Bardgett and van der Putten 2014; Wall et al. 2015), such as nutrient cycling, climate regulation, and disease and pest management (FAO et al., 2020). Earthworms (Annelida: Clitellata: Crassiclitellata) are a well-known member of soil macrofauna communities, and although they make up a very small proportion of species in the soil (Anthony et al. 2023), they are key players driving many ecosystem functions and services (Blouin et al. 2013). For example, they increase aboveground yield (van Groenigen et al. 2014), nutrient cycling rates, and decomposition (Huang et al. 2020).

IUCN Red List assessments are a common approach to assessing which organisms are at risk and which threats they are facing. The IUCN Red List protocol was created in 1994, providing a more quantitative framework to the assessment protocol that preceded it (Mace et al. 2008). Shortly after this development a framework for sub-global assessments was proposed, and Regional Red List assessments were adopted in 2003 (Mace et al. 2008). For both the Global and Regional Red List assessments, experts work using a standardised protocol (Rodrigues et al. 2006; Mace et al. 2008) extracting all available information and data on a species to provide an assessment on the likelihood of extinction (Mace et al. 2008). Whilst a global assessment examines the status of a species across the entirety of its range, the Regional Red List only evaluates the species within a small region or country (IUCN 2012). In undertaking the assessment, all available information on the species is combined, providing a valuable resource that can easily be disseminated (Rodrigues et al. 2006). In addition, the assessment can be used as a baseline to measure future actions, identify potential conservation sites, guide management plans (Rodrigues et al. 2006; Betts et al. 2020), and with additional analysis, determine conservation priorities (Fitzpatrick et al. 2007).

Despite their importance, soil organisms are rarely considered in conservation policies or assessments at the global level (Phillips et al. 2017). There are an estimated 5,755 described species/subspecies of earthworms across the globe (Brown et al. 2023; Misirlio□lu et al. 2023), but only ∼6% of them (361 species/subspecies) have been assessed in the Global IUCN Red List (Accessed on 8 February 2024; Table 1). Taxonomic biases within the Global IUCN Assessments are well known (Rodrigues et al. 2006), and thus it has been highlighted that Annelida should be a priority for Red Listing (Gerlach et al. 2014). This lack of representation is also found within the Regional Red Lists that have been published. Although there are a few Regional Red Lists that include earthworms, such as the Red Lists for Germany (Lehmitz et al. 2016), the Russian Federation (Geraskina and Kuprin 2021), Brazil (MMA, 2022), New Zealand (Buckley et al. 2015) and Australia (Australian Government n.d.), or other soil organisms (e.g., oribatid mites: Napierała et al., 2018, fungi: Dahlberg et al., 2010), these are the exception.

**Table 1.** The assessment given to each of the 361 earthworm species that have been assessed in the Global IUCN Red List, as well as the biogeographic realm that the species are found in. Data accessed 8th February 2024 from IUCN (IUCN 2024)

| Region | Extinct | Critically Endangered | Endangered | Vulnerable | Near Threatened | Least Concern | Data Deficient | Total |
| --- | --- | --- | --- | --- | --- | --- | --- | --- |
| Antarctic |  |  |  |  |  | 1 |  | 1 |
| Australasia | 2 | 5 | 11 | 6 | 2 | 45 | 78 | 149 |
| Indomalaya |  |  |  |  | 1 | 1 | 1 | 3 |
| Palaearctic |  | 1 | 1 | 2 | 8 | 27 | 35 | 74 |
| Neartic |  |  | 1 | 2 | 1 |  | 1 | 5 |
| Afrotropic |  |  |  |  | 2 |  |  | 3 |
| Neotropic |  | 1 | 3 | 2 | 7 | 30 | 84 | 127 |
| Total | 2 | 7 | 16 | 12 | 21 | 104 | 199 | 361 |

In Canada, regional assessments on the risk of extinction of a particular species are conducted by the Committee on the Status of Endangered Wildlife in Canada (COSEWIC), the framework of which is built upon the IUCN Red Listing protocols. Such assessments are an important way to understand and therefore mitigate or minimise the threats biodiversity faces. COSEWIC has completed over 850 species at risk assessments to date, including many invertebrates (COSEWIC 2023). However, the assessments of invertebrate species have so far been limited to arthropods (predominantly Lepidoptera) and molluscs (predominantly freshwater mussels). Outside of a few terrestrial snails, no soil organisms have been assessed (Naujokaitis-Lewis et al. 2022) and there is also no indication that taxonomic biases within COSEWIC action plans are improving (Creighton and Bennett 2019).

In Canada, most research on earthworms has examined the spread and effects of invasive, non-native earthworms (Addison 2009), as they can have severe negative impacts on ecosystems such as causing increased decomposition of litter layers in forests, thus changing the structure of the understory (Frelich et al. 2019). However, there may be as many as eight native earthworm species in Canada (Addison 2009). Unfortunately, of these eight, due to a lack of certainty in the identification of specimens (e.g., *Bimastos lawrenceae*) and the lack of knowledge of whether some widespread species were historically present (e.g., *Bimastos parvus*), only five species can be considered native with certainty (Reynolds 2022).

Here we present the results of a Regional Red List assessment performed for five native earthworm species found in Canada. By doing these assessments we aim to demonstrate that they are also feasible for understudied invertebrates. In addition, by collating all the information into the assessment, it can be used for setting conservation priorities based on earthworms (and by extension, for other soil taxa). Ultimately, our aim is to increase conservation focus towards life belowground.

## Methods

Of the eight earthworm species that are thought to be native to Canada, the taxonomic experts among us (GGB, SJ, JWR, EKC, JM) determined which were to be assessed. Although members of the genus *Bimastos* are present in Canada, it is less clear whether they are native to the country and therefore they were not included in the assessments (Table 2). The assessments of the remaining five species were done following all the guidelines provided by the IUCN for the Global and Regional Red List Assessments (IUCN 2012; IUCN Standards and Petitions Committee 2022).

**Table 2.** Eight earthworm species found in Canada. For the five species that the taxonomic experts deemed to be native to the country, results from the Regional Red List Assessments are given. Results include the areas that the species was found at, the estimated EOO within Canada (* indicates that, due to too few sites within Canada, the EOO was calculated using the sites from the USA), the main threats that the species faces, and the final assessment. No species listed in this table has been assessed as part of the IUCN Global Red List Assessment.

| Assessed Species | Canadian Location(s) | Estimated EOO in Canada (km<sup>2</sup>) | Main threats | Regional Assessment |
| --- | --- | --- | --- | --- |
| *Arctiostrotus fontinalis* | Yukon | 237,138* | - Residential & commercial development<br>- Biological resource use | Data Deficient |
| *Arctiostrotus perrieri* | Vancouver Island, Haida Gwaii, mainland coastal regions of southern BC | 44,080 | - Residential & commercial development<br>- Agriculture & aquaculture<br>- Natural system modifications<br>- Biological resource use<br>- Pollution<br>- Climate change & severe weather | Near Threatened |
| *Arctiostrotus vancouverensis* | Vancouver Island | 5,221 | - Residential & commercial development<br>- Agriculture & aquaculture<br>- Natural system modifications<br>- Biological resource use<br>- Pollution<br>- Climate change & severe weather | Vulnerable |
| *Sparganophilus tamesis* | New Brunswick, Quebec and Ontario | 256,991 | - Natural system modification<br>- Climate change & severe weather | Least Concern |
| *Toutellus oregonensis* | Vancouver Island | 2,595* | - Residential & commercial development<br>- Agriculture & aquaculture<br>- Natural system modifications<br>- Biological resource use<br>- Pollution<br>- Climate change & severe weather | Data Deficient |
| **Non-assessed species** | **General Remarks** |  |  |  |
| *Bimastos parvus* | Widespread and cosmopolitan species, even outside of North America (Csuzdi et al. 2017). |  |  |  |
| *Bimastos beddardi* | Potential synonym for *Bimastos parvus* (Csuzdi et al. 2017). Only found at two sites in Canada (Quebec) with a limited number of specimens, but unlikely to be an established species (Reynolds 2010). |  |  |  |
| *Bimastos lawrenceae* | Only 5 specimens, across two dates (1985 and 1993), found at one site in British Columbia (Douglas Peak, Vancouver Island) (McKey-Fender et al. 1994). |  |  |  |

Literature searches were conducted for *Arctiostrotus fontinalis, Arctiostrotus perrieri, Arctiostrotus vancouverensis* and *Toutellus oregonensis* (Family: Megascolecidae) and *Sparganophilus tamesis* (Family: Sparganophilidae) (Figure 1). Web of Science and Google (through Publish or Perish; Harzing, 2007) were searched using the keywords “[species binomial] + Canada”. After removing duplicate articles from the two searches, each article was reviewed and relevant information extracted into standardised templates. The standardised templates allowed the compilation of information relating to the taxonomy of the species, the geographical distribution using occurrence records (including exact locations of where the earthworms had been collected in Canada), population sizes, habitat types where the species had been found, as well as any threats the species face and which conservation actions were in place. The standardised template was designed to correspond with the IUCN assessment template.

**Figure 1.**
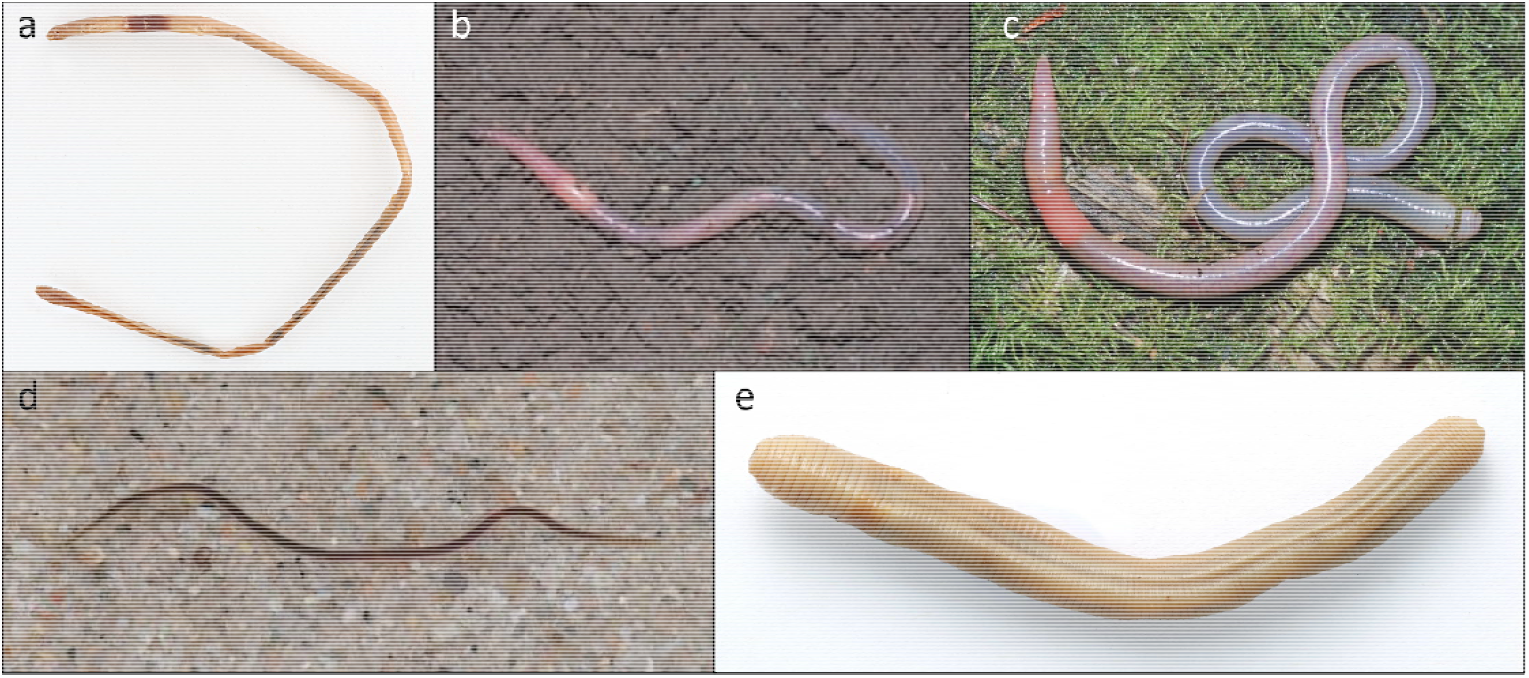
The five earthworms native to Canada that were assessed using the IUCN-based Red List Assessment. (a) *Arctiostrotus fontinalis*, (b) *Arctiostrotus perrieri*, (c*) Arctiostrotus vancouverensis*, (d) *Sparganophilus tamesis* and (e) *Toutellus oregonensis* (photographs from Reynolds, 2022, with permission)

For each species, the Extent of Occurrence (EOO) was calculated using the sites where each species had been found. All site coordinates were converted to decimal degrees, and if a site lacked coordinates but a location name was presented, this was matched to appropriate coordinates using Google Maps. A convex hull was then created around all points (i.e., the smallest polygon that could enclose all the sites), and the area of the convex hull was calculated as the EOO. Given the lack of population data, location data or habitat data (with the last two as defined by the IUCN) for the five species, the EOO, together with information on trends of threats, was the only criterion used for the assessments (discussed further below).

Across three meetings in 2021 and one meeting in 2022, the assessors (six of the authors of this paper) met to assess the five species. Information collated in the standardised data templates was reviewed, alongside the EOO maps. Information was fully compiled and entered into the assessment templates. The IUCN Red List category for each species was only determined following the entire review.

If there was insufficient data to make a full assessment, the species were marked as Data Deficient (DD). Lack of data typically corresponded to species that had only been found in a couple of locations within Canada, and thus evaluation based on the criteria below was not possible. For species that had sufficient data, the assessment could result in one of three threatened categories (Critically Endangered [CE], Endangered [EN], Vulnerable [VU]) or one of two categories of lower risk (Near Threatened [NT], Least Concern [LC]). Species are assessed as LC if they are widespread and abundant, and thus are not threatened, but are assessed as NT if they are close to qualifying for the threatened categories. As mentioned previously, the lack of population data for the five earthworm species meant that they were primarily assessed based on their EOO (criterion B1 of the assessment and associated sub-criteria). In brief, species can be classified as VU with an EOO < 20,000km^2^, EN with an EOO < 5,000km^2^, and CE with an EOO < 100km^2^. However, in order to meet the criteria of B1, the species also need to meet at least two of three sub-criteria: (a) severely fragmented or known to exist in few locations, (b) continuing decline, or (c) extreme fluctuation. Sub-criteria (b) and (c) can be represented by declines or fluctuations in any of the following: (i) extent of occurrence; (ii) area of occupancy; (iii) number of locations or subpopulations; or (iv) number of mature individuals (see IUCN guidelines for full details; IUCN 2012; IUCN Standards and Petitions Committee 2022). However, when one sub-criterion is met, but it is uncertain whether the second sub-criterion is met, the assessment category can be given as a range, with an indication of the most plausible category (IUCN Standards and Petitions Committee 2022).

Whilst the assessment was primarily completed using the EOO and therefore Criterion B, any information on population trends, habitat requirements, ecology, threats and conservation actions were compiled into the standardised templates, and where applicable, was used as additional evidence for the assigned assessment category.

## Results

The five species assessed belong to three genera: *Arctiostrotus, Sparganophilus* and *Toutellus. Arctiostrotus* and *Toutellus* are members of the Megascolecidae and are found only in Canada and the USA. *Sparganophilu*s is a member of the Sparganophilidae, and the species assessed here, *S. tamesis* (formerly *S. eiseni*), has been reported from Canada, USA, Mexico and several European countries. There are 12 additional species and two subspecies in the genus, but currently they are not known from Canada, only from the USA (Table 2).

Due to lack of data, two of the five species were classified as DD (*A. fontinalis* and *T. oregonensis*)(Phillips et al. 2022). Both species are known from only two sites within Canada, with only a few individuals found in total, although both species are also present in the USA (Figure 2a and d). Interestingly, in the case of *A. fontinalis*, there is considerable difference in habitats occupied by the Canadian population versus the US population. The Canadian locations were situated at >1,000 m above-sea level, whereas the USA sites were near sea level, implying three possibilities, either broad habitat requirements, habitat requirements that differ between subpopulations, or the individuals were different (sub-)species. Further investigation, potentially including DNA-based identification, would need to be conducted to ascertain.

**Figure 2.**
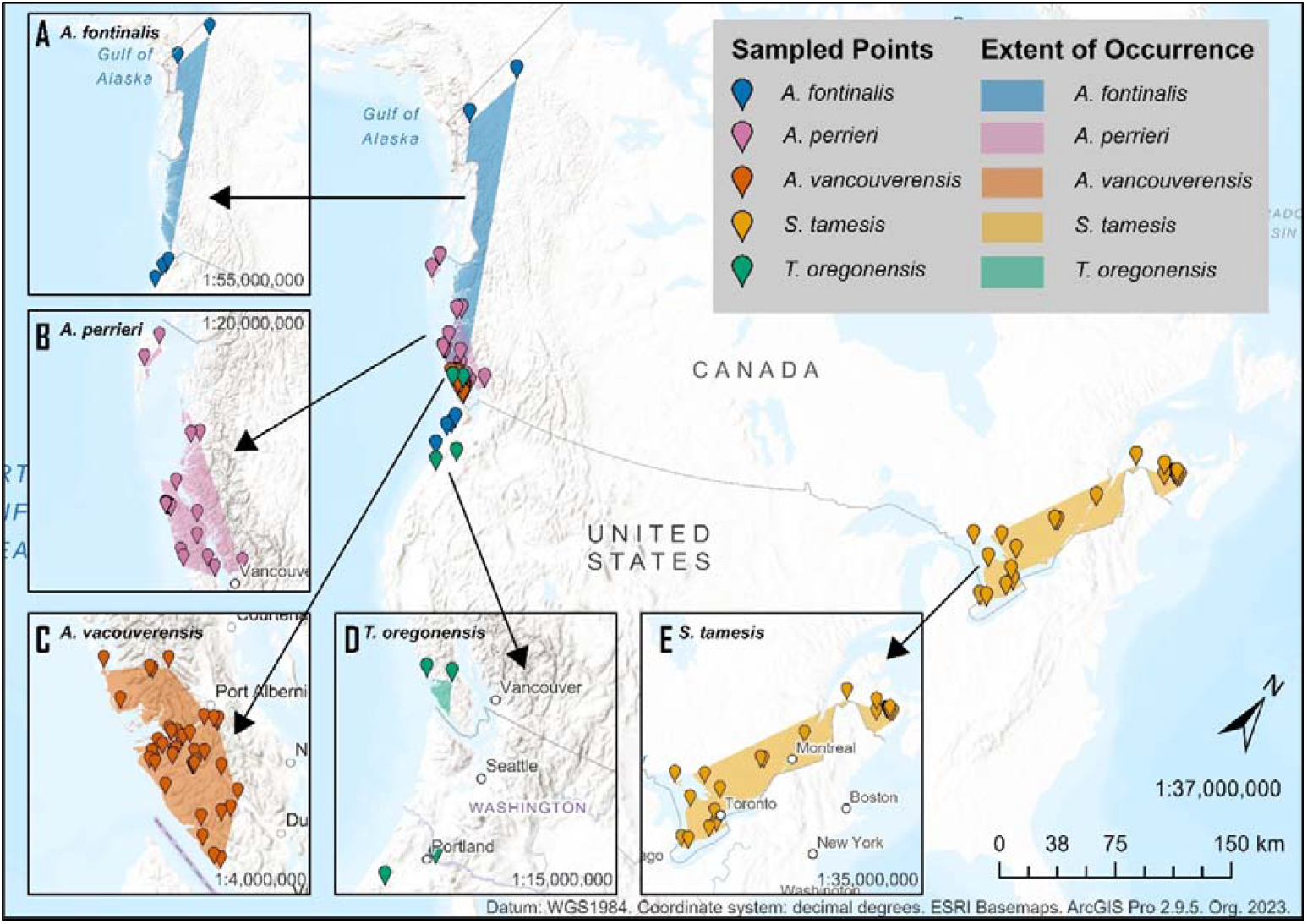
Main map indicates the sites where the five earthworm species have been recorded in Canada, as well as occurrences in the USA for *A. fontinalis* and *T. oregonensis*. Inset maps A-E show in greater detail the EOO within Canada for each of the species

Two species were not classified in the threatened categories (Phillips et al. 2022). *Sparganophilus tamesis* was classified as LC, due to its very large range across the Atlantic Maritimes (Figure 2e). *Arctiostrotus perrieri* was classified as NT, as although it has a large EOO (56,401 km^2^), it is only found on Vancouver Island and Haida Gwaii (Figure 2b). It is therefore subjected to a number of anthropogenic threats across the entirety of its known range, including an increase in urbanisation, climate change (increases in area, intensity and duration of forest fires, as well as increase in extreme weather events) and forestry activities.

*Arctiostrotus vancouverensis* was the only species to be classified within the threatened categories (Figure 2c; Phillips et al. 2022). The small EOO (5,221 km^2^) put the species well within the VU category. However, in order to be classified in a threatened category using the B1 criteria, two sub-criteria also need to be met. Given the lack of data on the number of locations (as defined by the IUCN), Criterion B1a could not be determined.

Criterion B1c also could not be determined, as the species has not been reported since 1994 (Marshall and Fender 2007), and therefore there is no way to determine whether there were extreme fluctuations in extent of occurrence. However, given the logging activities that occur within its EOO, as well as climate extremes (fires, floods, average increases in temperature and heat waves), it is likely that this species undergoes extreme population and/or EOO fluctuations as a result of the increase in mortality from these direct disturbances.

Criterion B1b was the only sub-criterion to be met, as due to a 35% increase in the number of dwellings being built on Vancouver Island between 2001 and 2021, as well as the increase in human population by 30% during the same timeframe (data from https://www.statcan.gc.ca/, accessed 12 September 2023), it can be strongly inferred that the EOO (Criterion B1b(i)) is declining as a result of increased disturbance and soil perturbations. Thus, *A. vancouverensis* was assessed as VU, but with a range of VU-NT, reflecting the uncertainty of not meeting both sub-criteria.

## Discussion

Using the Regional IUCN Red List Assessment, we were able to assess the extinction risk of five native earthworm species found in Canada; two as Data Deficient, and one each as Least Concern, Near Threatened and Vulnerable (Fig. 2, Table 2). Overall, despite our being able to classify three of the species in a non-Data Deficient category, there was little data available. Although some of the earthworm occurrence records dated from the 1930s (A. perrieri; McKey-Fender et al. 1994), there was considerably less research effort from more recent decades. We therefore call for increased awareness and more surveys of native species of earthworms across Canada, as well as of other soil organisms that could be potentially at risk (e.g., terrestrial snails; Nicolai and Ansart 2017). Without up-to-date distribution data it is not possible to fully understand the extinction risks of species, and the threats they face. We also hope that there will be more focus on soil organisms, and other invertebrates, by COSEWIC in the future.

The threats the earthworm species faced were very similar. All species but *A. fontinalis* are threatened by climate change. Unfortunately, Canadian greenhouse gas emissions have only declined by 1.1% between 2005 and 2019, despite pledges in the Paris Agreement to reduce greenhouse gas emissions by 30% by 2030 (Office of the Auditor General of Canada 2021). Thus, as a country, Canada is still contributing substantially to climate change. Given that the global average for the climate change velocity (the speed at which a species would need to move in order to stay within its climatic niche) is 0.42□km/year (Loarie et al. 2009), and earthworm populations have been estimated to spread at a rate of only 0.005 - 0.018 km/yr on their own (Marinissen and van den Bosch 1992; Cameron and Bayne 2014), it is unlikely that any of the species will be able to respond to the impacts of a changing climate via movement out of the unsuitable area without human assistance or a considerable shift in climate change trajectories.

The second most prevalent threat faced by all species except *S. tamesis* (the only native species in the east of the country) were factors related to urbanisation. Indeed, urbanisation is becoming an increasingly important threat to many species and their habitats (Li et al. 2022). It has been estimated that 93% of Canada’s endangered species are affected by habitat degradation, with the majority of the habitat degradation being a result of agricultural activity and urbanisation (Venter et al. 2006).

These five earthworm species face similar threats as other many species in Canada, and so conservation actions that are put in place for other species may benefit any of the earthworm species with overlapping distributions (Cameron et al. 2019). In 2021, 13.5% of terrestrial land in Canada was either conserved or protected (Environment and Climate Change Canada 2022) as a result of implementation actions towards the Aichi Target 13 (referred to as Canada Target 1 in Canadian policy). Of the ∼130 sites of earthworm occurrences across the five assessments, 15 sites were within a conservation area. Expansion of these areas, or creation of new protected and conservation areas, could benefit the conservation of earthworms where their range overlaps.

Due to the lack of data, two species had to be listed as DD in their assessments, *A. fontinalis* and *T. oregonensis*. However, if research efforts were put into gathering additional data, it is highly likely that those species would fall within the threatened categories. At the two sites these two species were found, very few individuals were collected, indicating that population sizes at these sites are likely to be very small. Additionally, previous studies have estimated that 56% of species listed as DD in the Global IUCN assessment may actually be threatened by extinction (Borgelt et al. 2022). Thus, not only are DD species of high conservation interest, they also often require the same level of conservation protection as species that have been assessed as threatened (Mace et al. 2008).

### Applicability of the IUCN Red List assessments for soil invertebrates

We have shown that it is possible to undertake IUCN Red List Assessments for understudied invertebrate species, and by using the standardised quantitative protocol provided by the IUCN these assessments can be compared with other taxa (Collen et al. 2016). Some authors have stated that they do not think the IUCN Red List Assessments are appropriate for invertebrates (Cardoso et al. 2011; Adriaens et al. 2015; Napierała et al. 2018). They question whether the thresholds for the threatened categories using Criterion B are appropriate for small organisms (such as invertebrates) that typically have small ranges. Additionally, as invertebrates are typically undersampled, this may result in a reduced size of the EOO and thus an overestimation of extinction risk (Cardoso et al. 2011). However, others have found that EOO is relatively robust with reduced sampling effort (Marsh et al. 2023). While noting that Criterion B is the most commonly used for invertebrates (Cardoso et al. 2011), it is also the most commonly used across the Global IUCN Red List as a whole (Collen et al. 2016). Additionally, it is difficult to assess the number of individuals in invertebrate populations, and assessors are therefore unable to assess how the population is changing over time (Criterion A)(Cardoso et al. 2011).

However, as others have countered, invertebrates are not unique with their issues in aligning with the different criteria, and problems exist in other taxa (e.g., fungi; Dahlberg and Mueller 2011; seaweeds; Brodie et al. 2023). As the assessments were developed for a broad range of species with diverse life histories (Mace et al. 2008; Collen et al. 2016) applying the assessment should not be discouraged.

In our assessments, particularly for *A. vancouverensis*, the most noticeably lacking data were related to number of ‘Locations’ of the species, and the habitat requirements of the species, both discussed further below. Having this information about the species would have allowed easier classification in the sub-criteria of Criterion B. Therefore, besides increasing our knowledge on the distribution of soil-dwelling invertebrates, we also call on researchers to further investigate ecological requirements of each species, as well as other information such as life history and population dynamics, which may help in future assessments.

The IUCN defines ‘Location’ as “*a geographically or ecologically distinct area in which a single threatening event can rapidly affect all individuals of the taxon present*.” (IUCN Standards and Petitions Committee 2022). While we know that *A. vancouverensis* is sensitive to increased temperatures (Fender 1995), we do not have accurate information on its exact tolerance limits, or data on other direct impacts from climate change that may be impacting the species now or in the future. Equally, the observations of the species are too sparse, including too temporally sparse, to match with data related to other threats (such as logging, or housing development) to calculate number of Locations using these other threats. Given the amount of data related to the threats that are available for Vancouver Island (and the wider BC area; https://catalogue.data.gov.bc.ca/), it was the lack of information about the connection between the threats and the species’ observations or its ecology that prevented us calculating the number of Locations. We are well aware that for many regions of the world and for many species globally, environmental data is not available. For understudied invertebrate species that may be relying on Criterion B for assessment, this poses even further problems.

As with Locations, the definition of ‘Habitat’ is also quite specific within the IUCN Assessments, specifically, *“[an] area, characterized by its abiotic and biotic properties…*..*avoid using generic classifications such as “forest” that indicate a biotope, a vegetation type, or a land-cover type, rather than a species-specific identification of habitat*” (IUCN Standards and Petitions Committee 2022). For more widely spread species, definition of the habitat type was possible; for example, *A. perrieri* is often found in mixed coniferous forests, such as spruce-hemlock (Spiers et al. 1986; McKey-Fender et al. 1994). However, for species with less distributional data, habitat could not be determined. For example, *A. vancouverensis* is known to be found in forested areas (Reynolds 2019), which is too general to match the requirements of the IUCN categories, and specifically found in humus and logs (McKey-Fender et al. 1994), a microhabitat that is unlikely to have data available for calculating habitat trends. Thus, ensuring that appropriate observations of the surrounding area are taken when collecting specimens, as well as reporting when species are *not* found in an area (especially within or near to the estimated range of the species), would help with determining habitat requirements.

### Incorporation of the Regional Red List assessments into national and global conservation frameworks

The species at risk assessments undertaken by COSEWIC are built upon the IUCN Red List assessment framework. Therefore, the information that we have gathered for the five earthworm species for these IUCN Red List Regional Assessments, and indeed any species assessed within the Red List framework, would be almost directly translatable to the COSEWIC database. The primary route for assessments by COSEWIC consists of three stages. Firstly, a list of possible species to assess is compiled by specialist subcommittees. Unfortunately, there is no specialist subcommittee for soil organisms, or even terrestrial invertebrates that are not arthropods or molluscs, which is potentially the main barrier for their inclusion in species at risk assessments. Secondly, all available data, knowledge and information is compiled. Finally, the species is assessed and results recorded. Although there is no relevant expert group for earthworms within COSEWIC, COSEWIC does accept unsolicited assessments (i.e., assessments created outside of the primary route), and whilst there is no guarantee that these unsolicited assessments would be accepted by COSEWIC, they seem the most viable option for getting soil organisms represented in the species at risk assessments in the short term.

As some of the species we assessed are endemic to Canada, the information that we have collated and their assessments would be directly applicable to the Global Red List, especially as none of the species are currently included. However, others, such as *S. tamesis*, would need additional information to be added as they are widespread outside of Canada (Rota et al. 2016). It is not uncommon for National Red Lists to include species not globally assessed. Brito et al., (2010) found that only 25% of the species on China’s National Red List had also been globally assessed. Similarly, Karam-Gemael et al., (2020) identified 596 Myriapoda species on Regional Red Lists around the world, and 210 species on the global red list; however, there were no species in common between any of the Regional Red Lists and the Global Red List. Given the typical lack of overlap between the two types of Red Lists (Dahlberg and Mueller 2011), this may provide some framework for prioritisation for the assessment, both global and regional, of soil biodiversity, and earthworms, in other countries. Undertaking regional assessments of soil-dwelling invertebrates that have already been globally assessed would provide additional understanding of the local threats they face, as well as identify potential conservation actions that can be done within a country.

## Conclusions

Despite earthworms being key players in many ecosystem functions and services, they are routinely omitted from conservation assessments, including IUCN Red List Assessments at global and sub-global levels. By undertaking regional assessments for five native earthworm species in Canada, we show that such assessments are feasible, despite rarity of locality and habitat data. These assessments can provide insights into where the species occur, the threats that they face and the knowledge gaps. Given that changes in climate, habitat loss and habitat degradation are increasing as a direct result of increases in human population, we call for more assessments of soil organisms to be undertaken and to be considered in future conservation efforts.

## Acknowledgments

This work was funded by a Canadian Institute of Ecology and Evolution working group grant and NSERC Discovery Grant (RGPIN-2019-05758 to EKC). In addition, HRPP received funding from European Union’s Horizon 2020 research and innovation program under the Marie Skłodowska-Curie grant agreement No. 101033214 (GloSoilBio). GGB received support from the Brazilian National Council for Scientific and Technological Development (Nos. 441930/2020-4 and 404191/2019-3). We thank Samantha Bennett, Jenacy Samways and Madison Silver for their help with extracting data from the literature. We also thank Marc W. Cadotte for his involvement with the working group.

## Statements and Declarations

This work was funded by a Canadian Institute of Ecology and Evolution working group grant and NSERC Discovery Grant (RGPIN-2019-05758 to EKC). In addition, HRPP received funding from European Union’s Horizon 2020 research and innovation program under the Marie Skłodowska-Curie grant agreement No. 101033214 (GloSoilBio). GGB received support from the Brazilian National Council for Scientific and Technological Development (Nos. 441930/2020-4 and 404191/2019-3). The authors have no relevant financial or non-financial interests to disclose. All authors contributed to the study conception and design. Data was collected by HRPP, and assessments were completed by GGB, SWJ, JM, JWR and EKC, and facilitated by HRPP. The first draft of the manuscript was written by HRPP and all authors commented on previous versions of the manuscript. All authors read and approved the final manuscript. The assessments created are available on Zenodo: https://doi.org/10.5281/zenodo.6625351.

